# Deletion of *proU* suppresses *proQ* phenotypes in *Escherichia coli*

**DOI:** 10.1101/2022.03.01.481631

**Authors:** Michelle N. Smith-Frieday, Craig H. Kerr, Janet M. Wood

**Author notes:** Address correspondence to: Janet M. Wood. MN Smith-Frieday, BWXT Canada, 581 Coronation Blvd, Cambridge ON, N1R5V3 and CH Kerr, Inceptive, 3160 Porter Dr, Palo Alto, CA 94304.

## Abstract

The ProQ protein interacts as an RNA chaperone with diverse RNA molecules in *Escherichia coli*. ProQ is implicated in the bacterial osmotic stress response. When the osmotic pressure is high, cells maintain their hydration by accumulating organic solutes denoted osmolytes. Transporters ProP and ProU (which is ProVWX) mediate osmolyte accumulation by *Escherichia coli*. Mutations at *proQ* impair ProP activity by reducing ProP levels (the ProQ transport phenotype) but do not impair ProU activity or reduce the level of ProX. The *proQ*^-^ bacteria are longer than *proQ*^+^ bacteria during growth in either low or high salinity medium and they grow slowly at high salinity (the ProQ growth phenotype). In addition, spherical cells with crescent-shaped, nucleic acid-rich foci appear and cells lyse (the ProQ morphological phenotypes). In this work, the *proQ* transport phenotype was suppressed by deletions of *proU*, or by an insertion of IS*5* in *proU*, when *proP* was expressed from the chromosome or from the heterologous, plasmid-based P_BAD_ promoter. A point mutation disrupting the Walker B motif of ProV inactivated ProU but did not suppress the transport phenotype. ProP activities and ProP levels varied in parallel, so *proQ* and *proU* act at the same level to regulate ProP expression. Deletion of the *proU* operon also suppressed the growth and morphological phenotypes. The *proU* locus may overlap the gene encoding a regulatory sRNA that acts with ProQ, contributing to cellular morphogenesis and osmotic stress tolerance, or the relationship between ProQ and *proU* may be indirect.

## INTRODUCTION

Non-coding RNAs and RNA-binding proteins are prominent participants in biological regulatory networks (1). Hfq and ProQ are global small RNA (sRNA) binding proteins in *Escherichia coli*. ProQ is a FinO-like protein that binds diverse RNA molecules and sRNA/mRNA complexes (2-5). The RNA-RNA interactomes of ProQ and Hfq overlap, and these proteins may compete for shared RNA targets (5). These observations and evidence that ProQ is an RNA chaperone (6) suggest that ProQ and Hfq are translational regulators of comparable significance (7). The structure of the ProQ protein is known (8, 9), and the mechanisms and physiological consequences of specific ProQ-RNA interactions are being explored (4, 10-14).

ProQ was first identified when mutations in the *proQ* locus conferred resistance to toxic proline analogue 3,4-dehydro-D,L-proline and decreased the activity of proline transporter ProP in *E. coli* (the ProQ transport phenotype) (15). Subsequent work showed that a Tn*5* insertion in *proQ* attenuated an osmotically induced increase in ProP activity (16). Osmotic stress perturbs cell structure, composition and function. Water flows out of cells as their medium becomes more concentrated (the osmotic pressure increases) and into cells as their medium is diluted (the osmotic pressure decreases). Bacteria attenuate such water fluxes by accumulating and releasing electrolytes (e.g. potassium glutamate) and small, uncharged or zwitterionic organic solutes called osmolytes (17). Cytoplasmic accumulation of osmolytes, mediated by transporters ProP, ProU, BetT and BetU, stimulates the growth of *E. coli* in high osmotic pressure media. Proton symporter ProP and ATP-binding cassette (ABC) transporter ProU (ProVWX) have broad substrate specificities, transporting proline, glycine betaine and related compounds (17). ProP is concentrated at the poles of *E. coli* cells whereas ProW, the membrane-integral component of ProU, is not (18, 19).

The transcriptional and posttranslational regulation of ProP and ProU have been extensively investigated (17, 20). Osmotic upshifts (increasing osmotic pressure) activate transcription of *proP* and *proU* on a time scale of minutes (20) and osmolyte uptake via transporters ProP and ProU on a time scale that is too short to measure with existing techniques. Substrates of ProP and ProU do not induce transcription of *proP* or *proU*. Rather, their transcription is determined by the osmotic pressure and other growth conditions (21). The *proP* locus is expressed and ProP can be activated in bacteria cultivated at low osmotic pressure whereas *proU* expression is minimal without osmotic induction (22). Osmotically induced potassium glutamate accumulation triggers global transcriptional responses to increasing osmotic pressure (23) and may activate transcription of *proP* and *proU* (24-26). Exogenous osmolyte glycine betaine decreases the expression of *proP* and *proU* in response to osmotic stimuli (21, 27-30). Cellular rehydration accompanies glycine betaine accumulation and may attenuate the signal that activates transcription (31). Despite their effects on ProP, *proQ* lesions do not alter ProU activity or ProX levels (32).

Early analyses suggested that ProQ affected ProP post-translationally (16, 33). Then ProQ was shown to be an RNA chaperone with an N-terminal FinO-like domain that has a higher affinity for duplex than single stranded RNA, a linker, and a C-terminal chaperone domain (6). Some small RNAs (sRNAs) work with a chaperone to bind the Shine-Dalgarno sequence within the 5’-untranslated region (5’ UTR) of a targeted mRNA and modulate translation (34). For example, FinO is a specific RNA chaperone that acts on FinP sRNA and a complementary sequence within the 5’-untranslated region (5’-UTR) of the *traJ* mRNA to accelerate its degradation. The impact of ProQ on ProP expression can be attributed to its N-terminal, FinO-like domain: plasmid-based expression of full length ProQ, of a ProQ variant without the linker, or of the N-terminal domain, raised ProP activity in bacteria with a *proQ* deletion (9). We therefore proposed that the FinO-like ProQ domain may act with an unidentified sRNA to regulate *proP* translation (6).

If ProQ followed a FinO-like mechanism, an antisense sRNA would interact with sequences in the *proP* 5’-UTR and/or the *proP* ORF. However, the *proP* 5’-UTR was not essential for the impact of ProQ on *proP* expression. Lesions in *proQ* attenuated ProP activity and decreased ProP levels when *proP* was expressed from a plasmid-based P_BAD_ promoter without the native *proP* 5’-UTR but retaining at least part of the *proP* 3’-UTR (6). Since sRNAs are also transcribed from intragenic regions or 3’-UTRs (35) it remains possible that an sRNA encoded within *proP* could act with ProQ to regulate *proP* translation. However, no such sRNA has been identified.

RpoD- and RpoS-dependent transcription from *proP* promoters P1- and P2-begins 182 and 95 base pairs upstream of the *proP* start codon, respectively (36, 37). Multiple post-transcriptional regulatory mechanisms elevate RpoS levels in osmotically stressed bacteria (38), indirectly affecting *proP* transcription from promoter P2. However, an *rpoS* defect did not significantly alter the impact of a *proQ* defect on *proP* expression, so ProQ does not act on *proP* via RpoS (6).

Recent work revealed that *proQ* lesions elongate *E. coli* cells, slow the growth of *E. coli* populations and alter cell morphology during cultivation in high salinity media (the *proQ* growth and morphological phenotypes) (32). Different aspects of these phenotypes could be associated with loss of ProQ or of Prc, an ATP-dependent periplasmic protease. The *proQ* locus is separated from the downstream *prc* locus by only 20 nucleotides, and evidence suggests that *proQ* and *prc* are co-transcribed (32, 39). Bacteria lacking ProQ but not Prc were elongated unless osmolyte glycine betaine was provided (32). Conversely, bacteria lacking Prc but not ProQ contained lysis-prone, spherical cells with an enlarged periplasm and an eccentric nucleus (32). Thus ProQ plays a role in cell morphogenesis, and its impact on ProP may be indirect. Here we reveal that the *proQ* transport phenotype is suppressed by deletions of or an IS*5* insertion in *proU*, but not by a point mutation that inactivates transporter ProU. Deletion of *proU* also suppresses the growth and morphological phenotypes.

## RESULTS

### A *proU* deletion suppresses the *proQ* transport phenotype

In bacteria with a wild type chromosomal *proP* locus, lesions at *proQ* decrease ProP activity without altering its osmotic pressure-dependence (the *proQ* transport phenotype) (33). The same phenotype was seen when *proP* was expressed from plasmid pDC79 (Fig. 1A). However, the *proQ* phenotype was not evident when *proP* was expressed from plasmid pDC79 in *E. coli* strain WG350 (Fig. 1B). We determined the origin of this suppression and assessed whether the *proQ* morphological phenotype was also suppressed.

**Figure 1:**
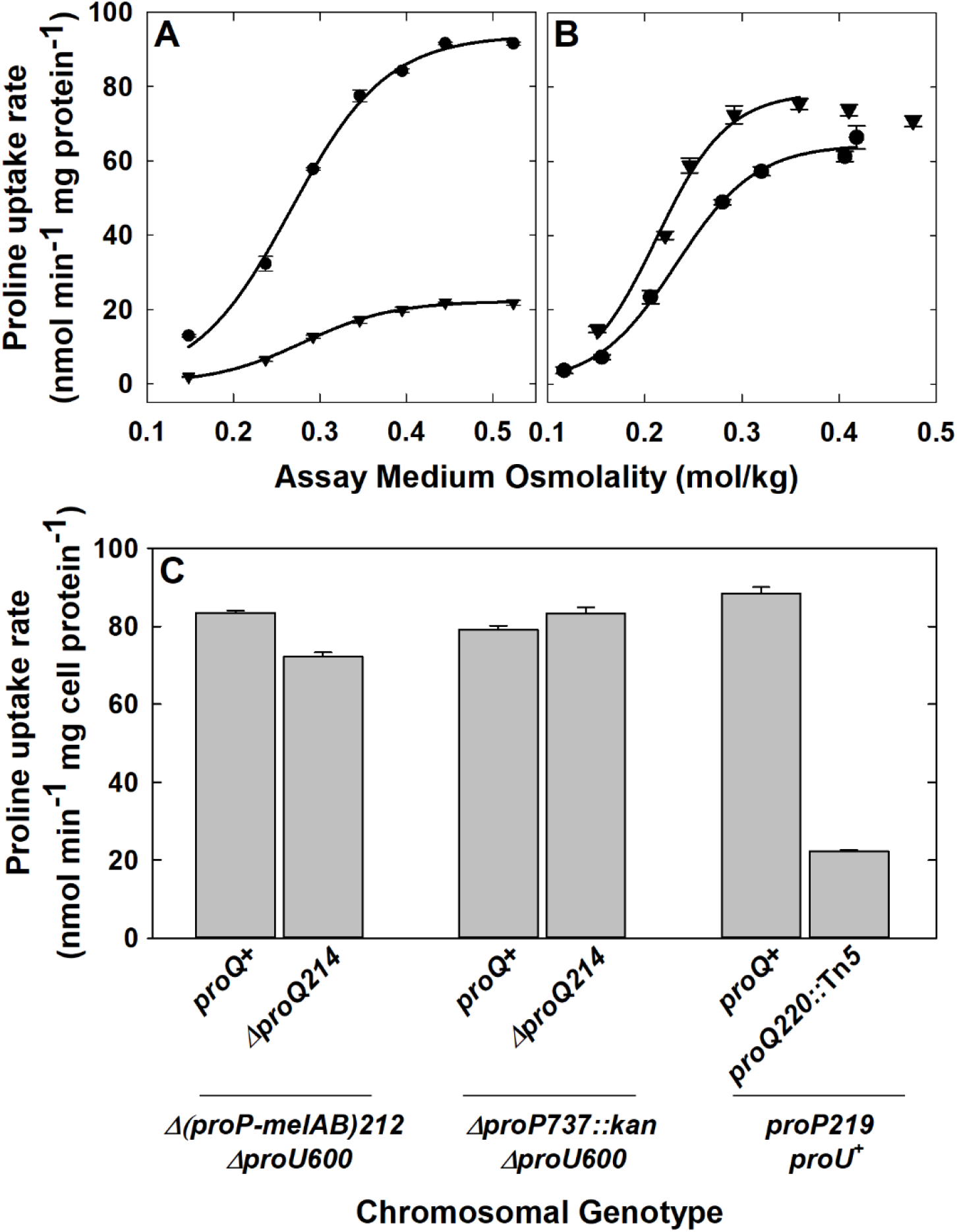
The *proQ* phenotype can be suppressed. Bacteria were cultured and and proline uptake activity was determined in MOPS media supplemented with NaCl to attain the indicated osmolalities as described in MATERIALS and METHODS. **A**: ProP activity in *E. coli* WG170 pDC79 (chromosome *proP219 proU*^*+*^) (circles) and its *proQ220::Tn5* derivative (WG1074 pDC79) (inverted triangles). The growth medium was supplemented with 50 mM NaCl (LOM, 0.25 mol/kg). **B**: ProP activity in WG350 pDC79 (chromosome *Δ(proP-melAB)212 ΔproU600*) (circles) and its Δ*proQ214* derivative (WG997 pDC79) (inverted triangles). The growth medium was NaCl-free (0.15 mol/kg). The *proQ* phenotype remained absent when the bacteria were cultured in MOPS media supplemented with NaCl to attain osmolalities in the range 0.15 through 0.72 mol/kg (not shown). **C**: ProP activity in *proQ*^+^ and *proQ*^-^ derivatives of strains WG350 pDC79 (chromosome *Δ(proP-melAB)212 ΔproU600*), WG1067 pDC79 (chromosome *ΔproP737::kan ΔproU600*) and WG170 pDC79 (chromosome *proP219 proU*^*+*^). In WG1067, *ΔproP737::kan* replaces *Δ(proP-melAB)212* so that the DNA segments flanking *proP* are restored but *proP* is not. The growth medium was supplemented with 50 mM NaCl (LOM, 0.25 mol/kg) and the assay medium osmolality was 0.47 mol/kg.

WG350 serves as a host for plasmid-based expression and analysis of proline transporters and their variants. It is devoid of proline uptake activity due to spontaneous deletions of proline transporter-encoding loci *putP, proP* and *proU* (40). ProP is routinely expressed in WG350 from plasmid pDC79 which was created by replacing a fragment of the multiple cloning site in vector pBAD24 (41) flanked by *Nco*I and HindIII sites with a DNA fragment extending from an NcoI site overlapping the *proP* initiation codon through a *Hind*III site 105 bp downstream of the *proP* termination codon (42). Thus *proP* expression from pDC79 is controlled by the vector’s P_BAD_ promoter, and the resulting *proP* transcripts lack the 5’UTRs present when *proP* is expressed from its native promoters (42).

The *proQ* phenotype was evident when the ProP activities of strains WG170 pDC79 and its *proQ220::Tn5* derivative were compared (Fig 1A). Thus pBAD24-based expression of ProP did not suppress the *proQ* phenotype. Different *proQ* lesions were present in the *proQ*^-^ derivatives of strains WG350 (WG997) and WG170 (WG1074): Δ*proQ214* and *proQ220::Tn5* respectively. (These and other relevant alleles are described in Table 2). However suppression of the *proQ* phenotype did not result from the difference in *proQ* genotype because both yielded the *proQ* phenotype in other contexts; various *proQ* insertion and deletion mutations affect *proP* activity in the same way ((33, 43), this study).

Strains WG350 and WG170 are both derived from *E. coli* RM2 which is *proP*^+^ *proU*^+^ (44, 45). WG350 harbours Δ*proU600* (46) and Δ(*proP-melAB)212* (40). The end points of these deletions are unknown, and potentially relevant genes, some with unknown functions, flank both loci. For example the loci flanking *proP* include upstream conserved genes of unknown function (*yjdN (*formerly *phnB* (47)*), yjdM (*formerly *phnA* (47)) and *yjcZ*) (48), upstream gene *yjdA* (which is now designated *crfC* (49)) and downstream genes encoding two-component system BasRS (50) and locus *eptA* (51) (Fig. 2). In contrast WG170 is wild type for *proU* and harbours a single base change in *proP* (Table 2). Using bacteria in which *proP* was expressed from plasmid pDC79, we tested the hypothesis that deletion of *proP, proU* or sequences flanking either locus suppressed the *proQ* phenotype.

**Figure 2:**
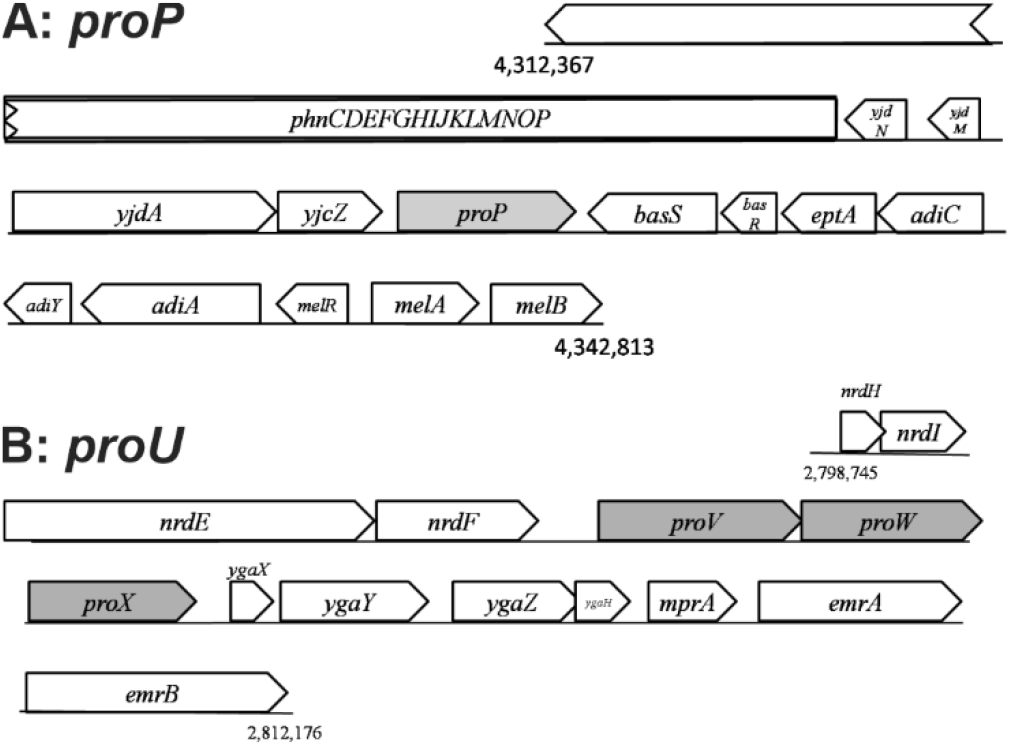
Contexts of the *proP* and *proU* loci in the *E. coli* chromosome. **A:** Loci surrounding *proP* from nucleotide (nt) 4,312,367 to nt 4,342,813. *yjdN* is also called *phnB* and *yjdM* is also called *phnA*. **B**: Loci surrounding *proU* (*proVWX*) from nt 2,798,745 to nt 2,812,176.

As noted above, the *proQ* phenotype was evident when plasmid pDC79 was introduced to *E. coli* WG170 and its *proQ220::*Tn*5* derivative (WG1074), both of which are *proP219 proU*^+^ (Fig. 1A and C). The *proQ* phenotype was not suppressed when each of the ORFs flanking *proP* was replaced in turn with a kanamycin resistance cassette in *E. coli* strains WG210 (*proP*^+^ *proQ*^+^ *proU205*) and WG914 (*proP*^+^ *ΔproQ676 proU205*) (Fig. 2 and 3). Strain WG210, which harbours a *proU* defect, was used for this analysis so ProP activity could be measured unambiguously. The impact of spontaneous mutation *proU205* is discussed further below. Furthermore, suppression remained when genes flanking *proP* were restored to strain WG350 and its Δ*proQ214* derivative (WG997) by introducing Keio collection allele *ΔproP737*::*kan* (Fig. 1C). These results suggested that suppression of the *proQ* phenotype did not result from deletion of the chromosomal *proP* locus or the flanking loci.

**Figure 3:**
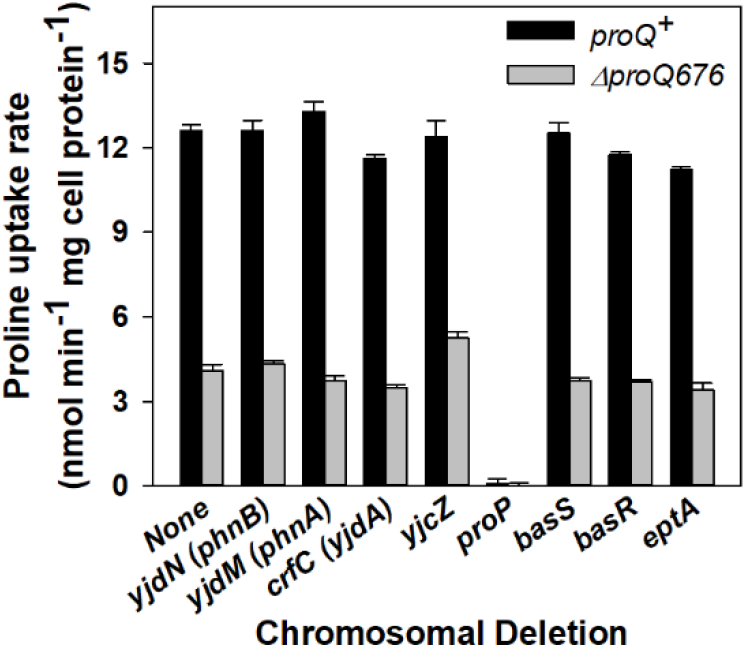
Effects of replacements of ORFs surrounding *proP* on ProP activity. Kanamycin resistance cassettes from the Keio collection (62) were introduced to strains WG210 (*proP*^*+*^ *proQ*^*+*^ *proU205 proQ*^+^) and WG914 (WG210 Δ*proQ676*), replacing *proP* and flanking ORFs. The loci replaced were *yjdN, yjdM, yjdA, yjcZ, proP, basS, basR* and *eptA*. Bacteria were cultured and ProP activity was measured in LOM as described in MATERIALS and METHODS.

Next the entire *proU* operon, from cryptic upstream promoter P1 through the *proX* termination codon, was deleted from strain RM2, creating strain WG1203 (*Δ(proV-proX)2098* (see MATERIALS and METHODS)). ProU and ProP confer high and moderate glycine betaine uptake activities, respectively, on *E. coli* cells cultivated and assayed at high osmotic pressure (MOPS medium supplemented with 300 mM NaCl) (22). As expected, deletion *Δ(proV-proX)2098* strongly decreased glycine betaine uptake activity under these conditions and the residual activity could be attributed to chromosomally encoded ProP (Fig. 4A). Western Blotting showed that it also eliminated ProX from this strain, further confirming deletion of the *proU* operon (Inset to Fig. 4A).

**Figure 4:**
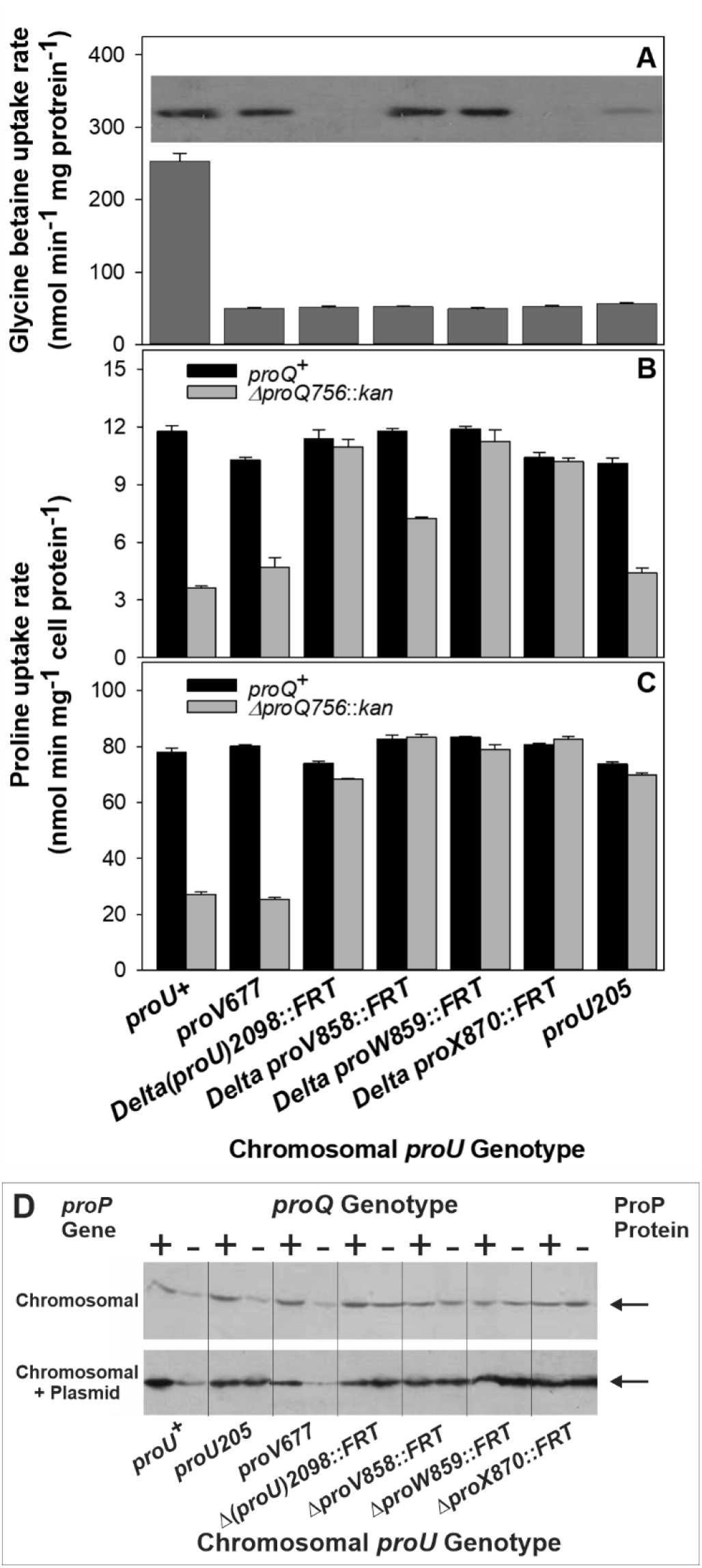
Effects of diverse *proU* defects on ProP and ProU. *E. coli* RM2 (*proP*^+^ *proU*^+^) and derivatives harbouring the listed *proU* alleles were cultured in MOPS medium. Glycine betaine (A) or proline (B,C) uptake activities were measured and ProX (A) or ProP (D) levels were determined by Western Blotting as described in MATERIALS and METHODS. **A**: **ProU activity and ProX levels** The growth medium was supplemented with 0.3 M NaCl to induce expression of the *proU* operon. Glycine betaine uptake activity was measured at an osmolality of 0.75 mol/kg as described in MATERIALS and METHODS. Both ProP and ProU contributed to the glycine betaine uptake activity of strain RM2 (*proU*^+^), whereas only ProP contributed to the activities of the other strains. *Inset*: The levels of the ProX protein in the bacteria employed for the transport assays were determined by Western blotting as described in MATERIALS and METHODS. **B-C**: **ProP activity** The growth medium was supplemented with 50 mM NaCl (LOM). Expression of *proU* is not induced under these conditions. Bacteria were *proQ*^*+*^ *or ΔproQ756*::*kan* (62). For C, the bacteria were transformed with plasmid pDC79 so *proP* was expressed from both the chromosome (the native *proP* locus) and the plasmid (under *araBAD* promoter control). Proline uptake activities were measured in NaCl-supplemented MOPS media with osmolalities of 0.46 mol/kg (B) or 0.25 mol/kg (C). **D**: **ProP levels** Western blot analysis of ProP levels in *proQ*^+^ and *proQ*^*-*^ variants (left and right lanes within each segment) of *E. coli* RM2 (*proU*^+^) and derivatives harbouring the indicated *proU* alleles. Bacteria were cultivated in HOM (top, 250 mM NaCl) or LOM (bottom, 50 mM NaCl) and *proP* was expressed from the chromosome (top) or both the chromosome and the plasmid (bottom). The *proQ* phenotype is indicated by higher ProP protein levels in extractcs from *proQ*^+^ than *proQ*^-^ bacteria (e.g. *proU*^+^ bacteria) and suppression of the *proQ* phenotype is indicated by similar ProP protein levels in extracts from *proQ*^+^ and *proQ*^-^ bacteria (e.g. bacteria with *proU* deletions).

No *proQ* phenotype was evident when allele Δ*proQ756::kan* was introduced to this strain (Fig. 4B, *Δ(proU)2098*::*FRT*), though the *proQ* phenotype was evident when strains RM2 and its Δ*proQ756::kan* derivative were tested as controls (Fig. 4B, *proU*^*+*^). Thus deletion of *proU* suppressed the *proQ* transport phenotype.

### The *proQ* transport phenotype is suppressed by deletions within *proU* but not by inactivation of ProU

Glycine betaine uptake via ProP or ProU lowers the expression of *proP* and *proU*. This effect has been attributed to attenuation of the signal activating their transcription (27, 28, 52). We assessed whether ProQ participates in this transport-dependent phenomenon by determining whether disruption of ProU function with minimal impact on the *proU* sequence would suppress the *proQ* phenotype. The *proU* disruption was designed as follows. ATP binding cassette protein MJ0769 is a component of a putative ABC transporter in *Methanococcus jannaschii*. Glutamate 171 immediately follows the Walker B motif in MJ0769, and replacement E171Q blocked ATP hydrolysis but not ATP binding or dimerization of MJ0769 (53, 54). To identify the corresponding residue in ProV, algorithm 3D PSSM (55) was used to align the predicted secondary structures of MJ0769 and ProV with the known secondary structure of MalK, the ATP binding protein of the maltose ABC transporter in *E. coli* (Protein Databank Identification Number 1Q1E). Among the ATP binding subunits of ABC transporters in *E. coli*, MalK was the subunit most similar in sequence to ProV for which a crystal structure was known. Residue E190 of ProV corresponded with E159 of MalK and E171 of MJ0769. Replacement E190Q was introduced to ProV by changing only 2 base pairs in the chromosomal *proV* locus of *E. coli* RM2 (yielding allele *proV677* in strain WG1199) (see MATERIALS and METHODS). Replacement E190Q abolished ProU activity as expected (Fig. 4A) but the *proU* operon was still expressed as the level of ProX was similar to that observed in *proU*^*+*^ bacteria (Fig. 4A *Inset*). Allele Δ*proQ756::kan* was introduced and ProP activity was measured, revealing that the *proQ* transport phenotype remained in the absence of ProU activity (that is, the *proQ* lesion impaired ProP activity, Fig. 4B). Thus the *proU* locus and not ProU activity is important for the *proQ* phenotype.

Next the impacts of in-frame deletions of *proV, proW* and *proX* were examined to localize the sequence within *proU* required for the *proQ* phenotype. As expected, each deletion eliminated ProU activity (Fig. 4A) and deletion of *proX* eliminated the ProX protein whereas deletion of *proV* or *proW* did not (Fig. 4A *Inset*). Allele Δ*proQ756::kan* was introduced and the proline uptake activities of the resulting strains were measured. The *proQ* phenotype was fully suppressed when *proW* or *proX* was deleted and partially suppressed when *proV* was deleted (Fig. 4B).

As noted above, the *proQ* transport phenotype was evident when *proP* was expressed from the chromosome of *E. coli* strain WG210 (Fig. 3 and 4B, *proU205*). WG210 harbours spontaneous mutation *proU205* (22). The nature of *proU205* was therefore determined (see MATERIALS and METHODS). This analysis revealed that IS*5* had inserted between the P347 and L348 codons of *proV*, abolishing ProU activity and decreasing but not eliminating *proX* expression (Fig. 4A).

Full or partial *proU* deletions influenced the *proQ* phenotype in the same way in bacteria without and with plasmid pDC79 (Fig. 4B and C). In contrast, insertion of IS5 in *proV* (*proU205*) did not suppress the *proQ* phenotype when *proP* was expressed only from the chromosome (Fig. 4B), but it did suppress when *proP* was also expressed from plasmid pDC79 (Fig. 4C). Deletion of *proV* (only, *ΔproV858*::*FRT*) partially suppressed when *proP* was expressed from the chromosome (Fig. 4B) and fully suppressed when *proP* was expressed from plasmid pDC79 (Fig 4C).

Data discussed above delineate the dependence of ProP activity on *proQ* and *proU*. The dependence of ProP protein levels on *proQ* and *proU* was also assessed by Western blotting (Fig. 4D). ProP activity and ProP protein levels varied in parallel (compare 4B with the top panel of 4D, and 4C with the bottom panel of 4D). Thus both *proQ* and sequences within *proU* regulate ProP activity by modulating ProP levels.

### Morphological effects of lesions in *proQ* and *proU*

Kerr *et al*. reported that *proQ*^-^ bacteria are longer than *proQ*^+^ bacteria during growth at either low or high salinity (the growth phenotype). In addition, the culture optical densities of *proQ*^-^ bacteria increase slowly at high salinity as spherical cells form and lyse (the morphological phenotype) (32). Like the *proQ* transport phenotype, most aspects of the *proQ* growth and morphological phenotypes were suppressed by a *proU* deletion (Fig. 5C) and not by a *proP* defect (Fig. 5B). The *proU* deletion partially suppressed cell elongation at high but not low salinity while fully suppressing cell rounding and lysis under both conditions. Thus, disruption of the relationships among *proQ, proP* and *proU* perturbs the rod shape and integrity of *E. coli* cells.

**Figure 5:**
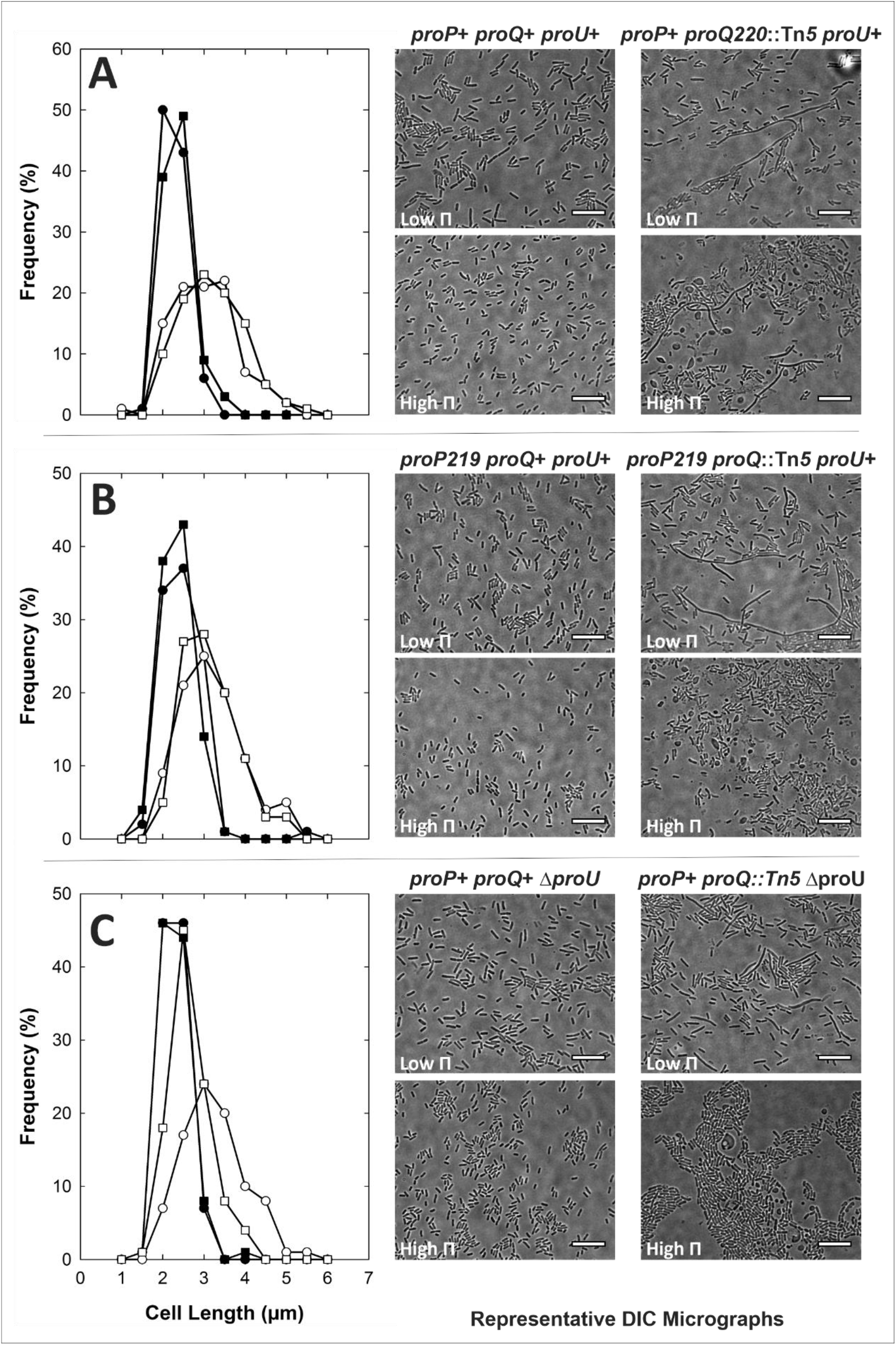
Impacts of *proP, proQ* and *proU* lesions on bacterial morphology. Bacteria were cultivated in MOPS medium to late exponential phase as described for transport assays (69) and visualized by DIC microscopy. Cell length distributions (left) and representative DIC micrographs (right) are shown for strains (A) RM2 (*closed symbols, proP*^+^ *proQ*^+^ *proU*^+^) and WG174 (*open symbols*, RM2 *proQ220*::Tn*5*), (B) WG170 (*closed symbols, proP219 proQ*^+^ *proU*^+^) and WG1074 (*open symbols*, WG170 *proQ220*::Tn*5*), and (C) WG1203 (*closed symbols, proP*^+^*proQ*^+^ *Δ(proU)2098*::FRT) and WG1327 (*open symbols*, WG1203 *proQ220*::Tn*5*). Growth media were supplemented with 50 mM NaCl (low osmotic pressure or Low Π (0.25 mol kg^-1^, *circles* in the left panels)) or 300 mM NaCl (high osmotic pressure or High Π (0.75 mol kg^-1^, *squares* in the left panels)). No *proQ*^+^ strain included cells with lengths greater than 6 µm whereas strains WG174 and WG1074 included cells with lengths greater than 6 μm at a frequency of 3-5% at both low and high osmolality. For strain WG1327 the frequencies were 12% and 0% at low and high osmolality, respectively. All scale bars correspond to 10 μm.

## MATERIALS AND METHODS

### Bacterial strains and plasmids

The relevant genotypes and immediate ancestors of the *E. coli* strains used for this study are listed in Table 1. Each was derived from strain RM2 (F^-^ *trp lacZ rpsL thi ΔputPA(101)*) (56), WG350 (RM2 *Δ(proP-melAB)212 Δ(proU)600*) (40), or BW25113 (F^-^ *Δ(araD-araB)567 ΔlacZ4787*(::rrnB-3) *λ*^*-*^ *rph-1 Δ(rhaD-rhaB)568 hsdR514*) (57). Plasmid pDC79 was created by replacing a fragment of vector pBAD24 (41) flanked by *Nco*I and HindIII restriction sites with a DNA fragment extending from an NcoI site overlapping the *proP* initiation codon through a *Hind*III site 105 bp downstream of the *proP* termination codon (42). Table 2 lists *proP, proQ* and *proU* alleles relevant to this study.

**Table 1:** Origins and relevant genotypes of *E. coli* strains used for this study^a^.

| Strain Name | From | Locus |  |  | Source or Derivation |
| --- | --- | --- | --- | --- | --- |
|  |  | *proP* | *proU* | *proQ* |  |
| JW5300 | BW25113 | *proP*<sup>+</sup> | *proU*<sup>+</sup> | $\Delta proQ756::kan$ | (62) |
| RM2 | CSH4 | *proP*<sup>+</sup> | *proU*<sup>+</sup> | *proQ*<sup>+</sup> | (56) |
| WG170 | RM2 | *proP219* | *proU*<sup>+</sup> | *proQ*<sup>+</sup> | (15) |
| WG174 | RM2 | *proP*<sup>+</sup> | *proU*<sup>+</sup> | *proQ220::Tn5* | (15) |
| WG210 | RM2 | *proP*<sup>+</sup> | *proU205* | *proQ*<sup>+</sup> | (22) |
| WG350 | RM2 | $\Delta(proP-melAB)212$ | $\Delta(proU)600$ | *proQ*<sup>+</sup> | (40) |
| WG914 | WG210 | *proP*<sup>+</sup> | *proU205* | $\Delta proQ676$ | (43) |
| WG997 | WG350 | $\Delta(proP-melAB)212$ | $\Delta(proU)600$ | $\Delta proQ214$ | this study |
| WG1063 | WG210 | $\Delta proP737::kan$ | *proU205* | *proQ*<sup>+</sup> | this study |
| WG1065 | WG914 | $\Delta proP737::kan$ | *proU205* | $\Delta proQ676$ | this study |
| WG1067 | WG350 | $\Delta proP737::kan$ | $\Delta(proU)600$ | *proQ*<sup>+</sup> | this study |
| WG1069 | WG997 | $\Delta proP737::kan$ | $\Delta(proU)600$ | $\Delta proQ214$ | this study |
| WG1072 | RM2 | *proP*<sup>+</sup> | *proU*<sup>+</sup> | $\Delta proQ756::kan$ | this study |
| WG1074 | WG170 | *proP219* | *proU*<sup>+</sup> | *proQ220::Tn5* | this study |
| WG1077 | RM2 | *proP*<sup>+</sup> | $\Delta proV758::kan$ | *proQ*<sup>+</sup> | this study |
| WG1078 | WG1077 | *proP*<sup>+</sup> | $\Delta proV858::FRT$ | *proQ*<sup>+</sup> | this study |
| WG1080 | WG1078 | *proP*<sup>+</sup> | $\Delta proV858::FRT$ | $\Delta proQ756::kan$ | this study |
| WG1082 | RM2 | *proP*<sup>+</sup> | $\Delta proW759::kan$ | *proQ*<sup>+</sup> | this study |
| WG1083 | WG1082 | *proP*<sup>+</sup> | $\Delta proW859::FRT$ | *proQ*<sup>+</sup> | this study |
| WG1085 | WG1083 | *proP*<sup>+</sup> | $\Delta proW859::FRT$ | $\Delta proQ756::kan$ | this study |
| WG1087 | RM2 | *proP*<sup>+</sup> | $\Delta proX770::kan$ | *proQ*<sup>+</sup> | this study |
| WG1088 | WG1087 | *proP*<sup>+</sup> | $\Delta proX870::FRT$ | *proQ*<sup>+</sup> | this study |
| WG1090 | WG1088 | *proP*<sup>+</sup> | $\Delta proX870::FRT$ | $\Delta proQ756::kan$ | this study |
| WG1119 | RM2 | *proP*<sup>+</sup> | *proU*<sup>+</sup> | $\Delta proQ856::FRT$ | this study |
| WG1199 | RM2 | *proP*<sup>+</sup> | *proV677* | *proQ*<sup>+</sup> | this study |
| WG1200 | WG1199 | *proP*<sup>+</sup> | *proV677* | $\Delta proQ756::kan$ | this study |
| WG1202 | RM2 | *proP*<sup>+</sup> | $\Delta(proV-proX)::kan$ | *proQ*<sup>+</sup> | this study |
| WG1203 | WG1202 | *proP*<sup>+</sup> | $\Delta(proV-proX)2098::FRT$ | *proQ*<sup>+</sup> | this study |
| WG1204 | WG1203 | *proP*<sup>+</sup> | $\Delta(proV-proX)2098::FRT$ | $\Delta proQ756::kan$ | this study |
| WG1327 | WG1203 | *proP*<sup>+</sup> | $\Delta(proV-proX)2098::FRT$ | *proQ220::Tn5* | this study |

<sup>a</sup>All strains designated WG were derived from *E. coli* CSH4 (F<sup>-</sup> *trp lacZ rpsL thi*) (44, 45). RM2 is CSH4 $\Delta putPA101$ (56), WG170 is RM2 *proP219* (15) and WG350 was derived from RM2 as described by Culham *et al.* (40). *E. coli* BW25113 is F<sup>-</sup> $\Delta(araD-araB)567 \Delta lacZ4787(::rrnB-3)$ $\lambda^- rph-1 \Delta(rhaD-rhaB)568 hsdR514$ (57).

**Table 2:** *proP, proQ* and *proU* alleles relevant to this study.

| **Allele** | **Description** | **Reference** |
| --- | --- | --- |
| *proP219* | Single base change (G to A at nt 1226 of the *proP* ORF) truncates ProP at A408 | (15, 33) |
| $\Delta(proP-melAB)212$ | Deletion encompassing *proP-melAB* and an unknown number of up- and downstream genes | (40) |
| $\Delta proP737::kan$ | Replacement of *proP* by a kanamycin resistance cassette | this work |
| $\Delta proP837::FRT$ | In frame deletion of *proP* | this work |
| *proQ220::Tn5* | Tn5 insertion after nt A314 of the *proQ* ORF interrupts the codon for E105 of ProQ | (15, 33) |
| $\Delta proQ214$ | *proQ* ORF replaced by an ORF encoding MENKCLSLVDGSLIVRAEHLVF | this work |
| $\Delta proQ756::kan$ | Replacement of *proP* by a kanamycin resistance cassette | this work |
| $\Delta proQ856::FRT$ | In-frame deletion of *proQ* | this work |
| $\Delta proQ676$ | *proQ* ORF replaced by an ORF encoding MENQPKCLCLGMGL | (43) |
| $\Delta(proU)600$ | Uncharacterized deletion of the *proU* operon | (46) |
| *proU205* | IS5 inserted between the codons for ProV residues P347 and L348 | (22),<br>this work |
| $\Delta(proV-proX)2098::FRT$ | Deletion of the *proU* operon (P1 promoter through *proX* termination codon) | this work |
| *proV677* | Encodes ProV-E190Q | this work |
| $\Delta proV758::kan$ | Replacement of *proV* by a kanamycin resistance cassette | this work |
| $\Delta proV858::FRT$ | In-frame deletion of *proV* | this work |
| $\Delta proW759::kan$ | Replacement of *proW* by a kanamycin resistance cassette | this work |
| $\Delta proW859::FRT$ | In-frame deletion of *proW* | this work |
| $\Delta proX770::kan$ | Replacement of *proX* by a kanamycin resistance cassette | this work |
| $\Delta proX870::FRT$ | In-frame deletion of *proX* | this work |

Routine DNA manipulation, plasmid construction, electrophoresis, and transformation were carried out as described previously (58, 59). Plasmid isolation was performed using QIAprep Spin Miniprep Kits (Qiagen, Mississauga, ON). The polymerase chain reaction (PCR) was performed as described previously (60) using *Pfu* turbo polymerase (Invitrogen, Burlington, ON). Site directed mutagenesis was performed using the QuikChange method (Stratagene Inc.) as described (61). Oligonucleotides were purchased from Cortec DNA Services (Kingston, ON). The Molecular Biology Supercenter (University of Guelph, Guelph, ON) performed DNA sequencing to verify all constructs. Genetic loci were deleted by introducing the relevant kanamycin (Km) resistance cassette from the Keio collection (62) by P1 transduction, then deleting the cassette as described by Datsenko and Wanner (57). Transductions were performed with phage P1 *cml clr_100* or P1*vir* as described by Miller (63).

Strain WG997 was created by introducing allele *proQ214* to *E. coli* WG350 in the same way that allele *proQ676* was introduced to *E. coli* WG210 (43), except that the allelic replacement vector was first reconstructed (pMS20 replacing pRAC2) so that the replacement open reading frame (ORF) encoded the expected peptide: MENKCLSLVDGSLIVRAEHLVF.

Allele *proU205* was isolated by selecting spontaneous mutant WG210 from strain WG170. WG210 was resistant to L-azetidine-2-carboxylic acid at high osmotic pressure (22). The *proU* loci of strains RM2 (*proU*^+^) and WG210 (*proU205* (22)) were compared to characterize *proU205*. A region of the *proU* locus extending from A47 of *proV* through C758 of *proX* was amplified by PCR with chromosomal DNAs from RM2 and WG210 as templates, and using primers proV1 (5’-CATCCACAGCGAGCGTTCAG-3’) and proX2 5’-GGGAAGCCATAATACGCACC-3’). The amplicon obtained with WG210 DNA was approximately 1 Kb larger than the (expected) 3 Kb product obtained with RM2 DNA. The former was purified and subjected to restriction endonuclease digestion with enzymes *Pvu*I, *Nde*I and *Sal*I. A 1 Kb insert was present upstream of the expected *Sal*I site near the beginning of *proW*. Sequencing revealed insertion sequence 5 (IS*5*) between the codons for ProV residues P347 and L348.

A point mutation designed to replace the glutamate at position 190 of ProV with glutamine was introduced to *E. coli* RM2 to create strain WG1199 (RM2 *proV677*). The *proVW* segment of the *E. coli* chromosome was amplified by PCR, adding *Bam*HI and *Hind*III restriction sites at the 5’ and 3’ ends respectively. This amplicon was inserted into vector pQE80L to create plasmid pMS26. The desired mutation in *proV* (*proV677*) was introduced to pMS26 by site directed mutagenesis, creating plasmid pMS27. A *BamH*I-*Sal*I restriction fragment of pMS27 was inserted into allelic exchange vector pKO3 (64) to create plasmid pMS31. Allele *proV677* was introduced to *E. coli* RM2 by allelic exchange as described by Link *et al*. (64), creating *E. coli* WG1199. PCR amplification and sequencing confirmed the presence of the desired mutation in the chromosome of strain WG1199.

A chromosomal deletion of the entire *proU* operon, extending from the cryptic P1 promoter (250 bp upstream from the initiation codon of *proV*) through the termination codon of *proX* was introduced to *E. coli* RM2 via the allelic exchange procedure of Datsenko and Wanner (57), as follows. Three DNA fragments were sequentially inserted into vector pGEM7Z (Promega Corp.) to yield plasmid pMS30 in which 500 bp of chromosomal DNA upstream of the P1 promoter and 500 bp of chromosomal DNA downstream of the *proX* termination codon flanked a kanamycin (Km) resistance cassette. The inserted fragments were obtained by PCR amplification using Keio Collection strain JW5300 (BW25113 *ΔproQ756::kan*) as template. The entire insertion constructed in this way was amplified and introduced to strain RM2 pKD46 (57) by electroporation, yielding transformant WG1202 in which the Km resistance cassette replaced the *proU* operon. The Km resistance cassette was removed by transforming strain WG1202 with plasmid pCP20 and screening for Km sensitive isolates as described by Datsenko and Wanner (57). PCR amplification and sequencing verified that strain WG1203 (*Δ(proV-proX)2098::FRT*) lacked *proU* but retained the scar (FRT) resulting from deletion of the Km resistance cassette.

### Bacterial cultures

Bacteria were cultivated in Luria-Bertani (LB) medium (63) or in MOPS medium (65) with NH^4^Cl (9.5 mM) as nitrogen source and glycerol (0.4% v/v) as carbon source, tryptophan (245 μM) and thiamine (1 mg/mL) to meet auxotrophic requirements. Low Osmolality Medium (LOM) was MOPS medium supplemented with 50 mM NaCl. High Osmolality Medium (HOM) was MOPS medium supplemented with 250 mM NaCl. Ampicillin (100 μg/mL) was added to maintain plasmids. Medium osmolalities (Π/RT, where Π is the osmotic pressure, R is the Gas Constant and T is the temperature (ºK)) were adjusted with NaCl and measured with a Wescor vapour pressure osmometer (Wescor, Logan, UT, USA). Cultures were grown at 37°C in a rotary shaker at 200 RPM. Optical densities were monitored with a Bausch and Lomb Spectronic 88 spectrometer.

### SDS−polyacrylamide gel electrophoresis and Western blotting

SDS−PAGE was performed as described by Laemmli (66) with gels containing 12% (w/v) polyacrylamide and 0.9% (w/v) bis-acrylamide. Gels were stained with Gel-Code Blue (Pierce, Rockford, IL) according to the manufacturer’s instructions. Western blotting was performed to detect ProQ (33, 43), ProP (67) and ProX (68) as previously described. Western blots were visualized with ECL reagents (GE Healthcare, Baie d’Urfe, QC) according to the manufacturer’s instructions.

### Transport assays

To determine proline or glycine betaine uptake rates, bacteria were grown in MOPS based minimal medium adjusted with NaCl to achieve the indicated osmolalities. ProP activity was measured as described previously (69). ProU activity was measured in the same way, at an osmolality of 0.75 mol/kg, using [1-^14^C]-glycine betaine as the substrate (10 µM, 5 Ci/mol).

### Protein assays

Protein concentrations were determined using the BCA (bicinchoninic acid) assay (70) with reagents from Pierce, (Rockford, IL) according to the manufacturers’ instructions and with bovine serum albumin as standard.

## DISCUSSION

ProQ deficiency decreases ProP levels (6) and impairs ProP activity (15, 16, 33) in *E. coli*. The effects of ProQ on ProP levels did not depend on the promoter used for *proP* transcription or the length of the 5’-UTR. In addition, *proQ* lesions reduced ProP levels when *proP* was expressed from a plasmid and the transcript lacked the *proP* 5’-UTR entirely ((6), Fig. 1 and 4D). An *rpoS* defect did not significantly alter the impact of a *proQ* defect on *proP* expression (6). These and other observations (summarized below) suggest that the impact of *proQ* lesions on ProP in *E. coli* are indirect, and not exerted via RpoS. However, it remains possible that a ProQ-binding sRNA transcribed from the *proP* open reading frame or 3’-UTR regulates *proP* translation.

Here we show that deletions or an insertion, not a point mutation, within the *proU* operon suppress the *proQ* phenotype (Fig. 4B-C). Deletion of *proW* or *proX* suppressed the *proQ* phenotype under all conditions tested (Fig. 4B-D), while a *proV* deletion (*ΔproV858::FRT*) and an insertion within *proV* (*proU205*) suppressed the *proQ* phenotype fully when *proP* expression was plasmid-based and only slightly (*ΔproV858::FRT*) or not at all (*proU205*) when expression was chromosome-based (Fig. 4). These observations suggest that a ProQ-binding sRNA may be encoded by a sequence within the *proU* operon, but no such sequence is evident by inspection or has been found experimentally. This is not surprising since the regions of complementarity between sRNAs and their mRNa targets may be short and discontinuous (71). Synthesis of such an sRNA could depend on growth conditions that affect the structure of the chromosome in the *proU* region, competition for transcription between the sense *proU* mRNA and the antisense RNA, and association of H-NS with the *proU* NRE (extending from the *proU* 5’-UTR into *proV*). No survey of sRNAs present in *E. coli* under osmotic stress has been reported. Barnhill *et al*. identified a 154 base sRNA (sRNA2111563), induced during desiccation of *Salmonella enterica* sv Typhimurium, that was predicted to anneal with *proP* (14), and ProP contributes to the desiccation tolerance of *S. enterica* (72). However, this sRNA has not yet been shown to regulate *proP* expression and it is not transcribed from sequences within *proP, proU* or *proQ*. In sum, the location of the gene encoding the putative sRNA remains unknown and that sRNA may not associate with *proP* mRNA. In fact, the ProQ transport phenotype may be secondary to newly discovered impacts of *proQ* defects on bacterial cell structure.

Lesions in *proQ* have two distinct effects on the growth and morphology of *E. coli* cells (32). First, *proQ*^-^ bacteria are longer than *proQ*^+^ bacteria after growth at low or high salinity ((32) and Fig. 5A, B). This effect was partially suppressed by deletion of the *proU* operon for bacteria cultivated at high but not low salinity (Fig. 5C). Second, polar effects of *proQ* lesions on downstream locus *prc* cause the formation of lysis-prone spherical cells bounded by concentric membranes during cultivation at high salinity ((32) and Fig. 5). This phenomenon was fully suppressed by deletion of *proU* (Fig. 5C).

The elongation of *proQ*^-^ bacteria cannot be a consequence of decreased ProP activity since the phenomena illustrated in Fig. 5 occurred during bacterial growth in the absence of ProP or ProU substrates. (ProU activity was unaltered under these conditions.) It is therefore likely that the primary ProQ targets have yet to be identified and decreased ProP levels are secondary to morphological effects of *proQ*^-^ mutations. ProP concentrates with cardiolipin at the poles and near the septa of *E. coli* cells (18). The decreased ProP levels in *proQ*^-^ bacteria could reflect a decrease in proportion of cytoplasmic membrane located at cell poles, though no link between ProP localization and *proP* expression has been reported. Decreased ProP activity could also reflect effects of membrane lipid on ProP activity (73, 74).

In sum, data presented in this paper suggest that the impact of *proQ* mutations on ProP activity (the ProQ transport phenotype) is secondary to the *proQ* morphological phenotype. ProQ may participate in the regulation murein synthesis, ensuring cell integrity during growth at high salinity.

## ACKNOWLEDGEMENTS

We are grateful for Discovery Grants OPG0000508 and 508-2008, awarded to JMW by the Natural Sciences and Engineering Research Council of Canada, and an Ontario Graduate Scholarship in Science and Technology, awarded to MS-F.

